# Effects of post-training novelty exposure on contextual fear memory: An attempt to translate behavioral tagging to humans

**DOI:** 10.1101/479659

**Authors:** Mason McClay, Joseph E. Dunsmoor

**Affiliations:** University of Texas at Austin, Department of Psychiatry, Austin, TX 78701, USA

## Abstract

In rodents, poorly formed hippocampal memories can be improved by novelty exploration within a critical time window, in line with the “behavioral tagging” hypothesis. Here, we sought to establish an analogue protocol to investigate if novelty exploration similarly operates to rescue weak hippocampal-dependent memories in humans. Adult humans underwent suboptimal contextual fear conditioning, followed 10 minutes later by open field novelty exploration in immersive 3D virtual reality. Novelty exploration did not improve long-term contextual fear memory, contrary to a behavioral tagging hypothesis. Despite this null result, we suggest further attempts to translate behavioral tagging from rodents to humans is warranted.

---

Why are some experiences transformed into a long-term memory while others are forgotten? An emerging line of research in rodents shows that weakly formed memories can be stabilized by a different and more salient experience that involves a common neural substrate (Moncada and Viola 2007; Ballarini et al. 2009; Wang et al. 2010; de Carvalho Myskiw et al. 2013). Referred to as behavioral tagging, there is now abundant evidence that hippocampus-dependent learning tasks are better remembered if learning is accompanied by novelty exposure. In rodents, this includes tasks for spatial learning (Wang et al. 2010), object recognition (Ballarini et al. 2009), inhibitory avoidance (Moncada et al. 2011), and contextual fear conditioning and extinction (Ballarini et al. 2009; de Carvalho Myskiw et al. 2013; de Carvalho Myskiw et al. 2014; Menezes et al. 2015). In the behavioral tagging model, weak learning sets the conditions for a long-term memory; but in line with the synaptic tag-and-capture model (Frey and Morris 1997), long-term memory formation is dependent upon stronger synaptic potentiation within overlapping neural ensemble during a critical time window around the time of training (Nomoto et al. 2016). Novelty exploration provides the conditions for strong synaptic potentiation by triggering upregulation of plasticity related proteins in the hippocampus (Straube et al. 2003) necessary to stabilize local learning tags set by the weak learning experience. Behavioral tagging has received considerable interest as a model to explain how weakly learned experiences are nonetheless retained in memory (Rogerson et al. 2014; Fernández and Morris 2018). But evidence of behavioral tagging in humans is limited, and the memory tasks used in human research to date are quite dissimilar from those used in rodents (Ballarini et al. 2013; Dunsmoor et al. 2015). The goal of the present study was to establish a proof of concept for translating a behavioral tagging protocol from rodents to humans. We attempted to maintain and adapt key elements of a hippocampus-dependent learning task (contextual fear conditioning) and novel open field exploration from the animal learning literature to determine whether post-training novelty exploration can rescue suboptimal hippocampus-dependent learning in humans.

Laboratory animal research on behavioral tagging has made progress using tasks for which the neural circuitry of learning and memory are extremely well-detailed. Contextual fear conditioning is a popular paradigm for basic research on learning and memory across species and offers a number of advantages for translational research (Maren et al. 2013). Context conditioning involves learning the association between spatiotemporal cues (i.e., a context) and an aversive unconditional stimulus (US, such as an electrical shock). As compared to standard Pavlovian cued conditioning that involves a discrete conditional stimulus (CS, such as a tone), in context conditioning the environment itself (usually a cage in rodent research) acquires the capacity to elicit a conditional fear response. The neurobiology of contextual fear conditioning is increasingly well detailed and relies predominately on connections between the hippocampus and basolateral amygdala (Fanselow and Poulos 2005; Marek et al. 2018). Importantly, expression of a long-term contextual fear conditioning memory is susceptible to improvement by a different (and stronger) hippocampus-dependent experience within a critical time window around the time of training. For instance, a weak contextual fear conditioning protocol consisting of just a few shocks over a short time interval is either preceded or followed by exploration of a novel open field (Ballarini et al. 2009). Novelty exploration improved long-term memory of weak contextual fear conditioning, as compared to control groups who either underwent weak training without novelty exploration or weak training followed by exploration of a familiar environment.

The present study tested the behavioral tagging hypothesis using contextual fear conditioning in 40 healthy adult human volunteers. Unlike rodent studies, context conditioning in human subjects research is typically conducted using a discrimination protocol that includes a context associated with shock (CTX+) and a within-subjects control context that is always free of shocks (CTX-). The dependent measure of contextual fear conditioning typically involves heightened fear-potentiated eye-blink startle responses within the CTX+ as compared to the CTX- (Grillon 2008). Here, two groups of subjects first underwent a purposefully weak contextual fear conditioning protocol presented using commercially available immersive virtual reality hardware (see Kroes et al. 2017 for details on context conditioning in immersive VR). Subjects were passively guided through the CTX+ and CTX- four times each. On one of the CTX+ trials, they received one unsignaled shock; on a second CTX+ trial, they received two unsignaled shocks; on the other two CTX+ trials, and on all CTX- trials, they received no shocks. Each trial was 30 seconds, and always followed by an intertrial interval (ITI) that consisted of passive navigation down a virtual hallway without any shocks.

After a ten minute break following conditioning, subjects either underwent novelty exploration (N = 20) or were in a control group (N = 20). To approximate open field novelty exploration, subjects in the novelty exploration group played an immersive virtual reality video game (*Eagle Flight*, Ubisoft, Montreal, Canada) through the Oculus Rift head-mounted display that incorporates aspects of novelty, reward, exploration, and navigation. The control subjects, on the other hand, passively viewed a 2D movie of the video game. Subjects in both groups returned 24-hours later to test fear-potentiated startle in the CTX+ and CTX- (4 trials each) without any shocks. We sought to establish that a weak contextual fear conditioning protocol would nonetheless yield a stronger contextual fear memory if weak training was accompanied by novelty exploration the previous day. Further descriptions of the task are detailed toward the end of the manuscript.

Results from the fear-potentiated startle analysis of context conditioning (Day 1) confirmed poor discrimination learning between the CTX+ and CTX- in both groups, as intended. There were a total of 8 startle probes each in the CTX+, CTX-, and ITI. The first and second half of conditioning were separated to examine whether learning improved over time. Repeated measures ANOVA using context (CTX+, CTX-) and time (early, late) as within-subjects factors and group (Exploration, Control) as between-subjects factors showed no effect of context (*F*_1, 38_ = 1.708, *P* = .199), Group (*F*_1, 38_ = 1.030, *P* = .316), nor a context by Group interaction (*F*_1, 38_ = 2.294, *P* = .138). There was an effect of time (*F*_1, 38_ = 35.375, *P* < .001), but no interactions with time. During early conditioning, neither group showed elevated startle in the CTX+ or CTX- as compared to the ITI (all *P*s > .09); but by late conditioning startle was elevated in CTX+ and CTX- as compared to the ITI in both groups (all *P*s < .002). Thus subjects appeared to learn that the hallway was safe, but startle was undifferentiated within the two contexts, confirming poor initial learning during the context conditioning session on day 1.

Subjects returned 24 hours later to test for startle using the same contexts as before, but without any shocks. Again, startle was separated into the first and second half of testing to examine potential effects of time on threat expression during testing. Unlike during context conditioning, there was a main effect of context (*F*_1, 38_ = 13.135, *P* = .001, partial eta^2^ = .257), but no effect of Group, and no interaction with Group (*F*_1, 38_ = 0.011, *P* = .918, partial eta^2^ < .001). There was an effect of time (*F*_1, 38_ = 74.994, *P* < .001, partial eta^2^ = .664), but no interactions with time. Follow-up t-tests on startle discrimination between CTX+ and CTX- showed no difference in the Novelty Exploration group during early (P = .4) or late (P =.14) testing. Likewise, the Control group did not exhibit discrimination during early testing (P = .06); however, startle was different in these contexts during late testing (*t*_19_ = 3.680, *P* = .002) such that startle was elevated in late CTX+ versus CTX- trials. However, an independent samples t-test on the CTX+/CTX- difference score during late testing did not show a significant difference in discrimination performance between groups (*t*_38_ = 8.879, *P* = .385). Consequently, we cannot conclude that the Control group showed *better* startle discrimination than the Novelty Exploration group. During early testing, both groups showed elevated startle in the CTX+ as compared to the ITI (both *P*s < .015), but not the CTX- compared to the ITI (both *P*s > .18); during late testing startle was elevated in CTX+ and CTX- as compared to the ITI in both groups (all *P*s < .001). In sum, there was no evidence that novelty exploration 10 minutes after weak context conditioning subsequently improved long-term memory performance in humans.

The goal of the present study was to develop a protocol to translate behavioral tagging research from rodents (e.g., Moncada and Viola 2007; Ballarini et al. 2009; Wang et al. 2010; de Carvalho Myskiw et al. 2013; Nomoto et al. 2016; Gros and Wang 2018) to humans. To date, there is some indirect evidence in support of a behavioral tagging mechanism in human episodic memory research. For instance, Pavlovian conditioning to exemplars from an object category (e.g., pictures of animals) has been shown to selectively and retroactively enhance memory for different exemplars from that category encoded prior to conditioning (Dunsmoor et al. 2015; Patil et al. 2017). This retroactive enhancement in memory was only observed following a delay (not immediately). Also, this effect was not observed if the items were strongly encoded prior to conditioning, in line with the hypothesis that behavioral tagging is a mechanism for strengthening memory for weakly learned experiences, not experiences that were already firmly established. Behavioral tagging has also been invoked to explain how exposure to novelty improved memory in elementary school children for a story, if the story was followed by a novel and interactive science demonstration (Ballarini et al. 2013). And physical activity several hours after learning (but not immediately) has been shown to improve hippocampus-dependent associative memory on a cued-recall test (van Dongen et al. 2016). But the link between enhancements of episodic memory and a behavioral tagging mechanism in humans is indirect, and lacks a direct translational analogue to the growing body of rodent studies using different measures of hippocampal memory function.

Here, we used contextual conditioning because it is a widely used paradigm that easily translates across species, and has been used in a number of behavioral tagging studies in rodents. The advantage to a cross species protocol is the ability to test a variety of conditions that should be met in order to evaluate behavioral tagging. For instance, any effects consistent with behavioral tagging should be time-dependent such that novelty exploration should only improve memory if it occurs close in time to another hippocampus-dependent task. An enhancement in memory should also be observed only after a delay and not immediately. Here, we did not find evidence that temporally pairing contextual conditioning with rewarded novelty exploration in immersive 3D virtual reality improved long term memory expression beyond a control group, who only passively viewed the novel environment in 2D. This null result could call into question whether context conditioning is sensitive to behavioral tagging processes in humans. Alternatively, failure to see an effect of novelty exploration on long-term context conditioned threat memory could be a constraint in how contextual conditioning is evaluated in humans.

One methodological challenge in translating contextual conditioning protocols to humans is the use of a within-subjects control context (the CTX-), which are unusual in rodent context conditioning protocols. We had predicted improved contextual threat memory would be revealed by discrimination between the CTX+ and CTX- the next day; but it is possible that novelty exploration has effects on promoting memory generalization between the CTX+ and CTX-, explaining the lack of differential startle between contexts. However, this explanation is undercut somewhat by recent work showing that extended training on a 3D spatial video game improves behavioral pattern separation (Clemenson and Stark 2015), suggesting that hippocampal-training should promote better discrimination.

Although we did not find evidence for behavioral tagging in a contextual conditioning, future efforts to investigate behavioral tagging in humans is warranted. It may be valuable, for instance, to test short-term (e.g., on Day 1) and long-term memory within subjects. However, effects of behavioral tagging on implicit fear memories in humans may simply be challenging to observe; there are methodological factors (e.g., the aforementioned CTX-), as well as inter- and intra-individual variability in physiological measures and subjective shock intensity to consider. In humans, behavioral tagging might be more pronounced in tasks that involve explicit memory processes, or spatial navigation, rather than implicit fear conditioning. Future work should consider whether behavioral tagging can rescue weak spatial object learning in humans, which would still be in keeping with several rodent investigations of behavioral tagging (Ballarini et al. 2009; Wang et al. 2010; Nomoto et al. 2016; Moncada 2017).

Subjects were 40 healthy adults (29 females; mean age = 21.57 y/o; SD = 2.3 years; range = 19 to 27) randomly assigned to either the novelty exploration or control group (N = 20 each). Participants provided written informed consent and the study was approved by the University of Texas at Austin IRB. Electromyography of the orbicularis muscle of the right eye collected at 2000 Hz was used to index fear-potentiated startle. To induce a startle reflex, a 50 millisecond 100 dB white noise was presented binaurally through headphones. Prior to the start of the experiment on both days, subjects were habituated to the startle probe through 9 presentations of the probe. To quantify startle responses during the experiment, the baseline EMG response over a 500 ms period prior to the onset of the startle probe was subtracted from the maximum EMG response from a 20-120 millisecond window after the startle probe. These raw scores were transformed to T-scores (z-score*10 +50) in keeping with prior research incorporating fear-potentiated startle in humans. The 6 millisecond electrical shock was delivered to the right wrist using the BIOPAC STM200 module (BIOPAC Systems) calibrated to a level deemed highly annoying but not painful. Results from statistical analyses were considered significant at *P* < .05.

The experimental design is shown in Figure 1. The context conditioning protocol was a slight variation to a recently published task (Kroes et al. 2017), and some technical aspects of the VR context conditioning protocol and procedure are described in more detail in Kroes et al. The environments were designed in Unity (Unity Technologies) using C#. The script included TTL pulses to trigger presentation of the startle probes and shocks, and to record event timing for psychophysiological analysis. The environment consisted of two distinct rooms. One of the rooms was blue and included pictures of different animals on the wall; one of the rooms was yellow and included pictures of different tools on the wall. The color and category of pictures on the walls were used to help subjects discriminate between the two contexts (and thus prevent generalization), as is common in discriminative fear conditioning protocols. The two rooms were separated by a hallway with gray walls. Subjects were passively guided down the hallway between each trial; thus the hall served as the inter-trial interval (ITI). Assignment of the blue or yellow room as CTX+ and CTX- was counterbalanced between subjects. Prior to the start of conditioning, subjects were familiarized with the Oculus Rift headset and given 1 minute in each room to actively explore.

**Figure 1.**
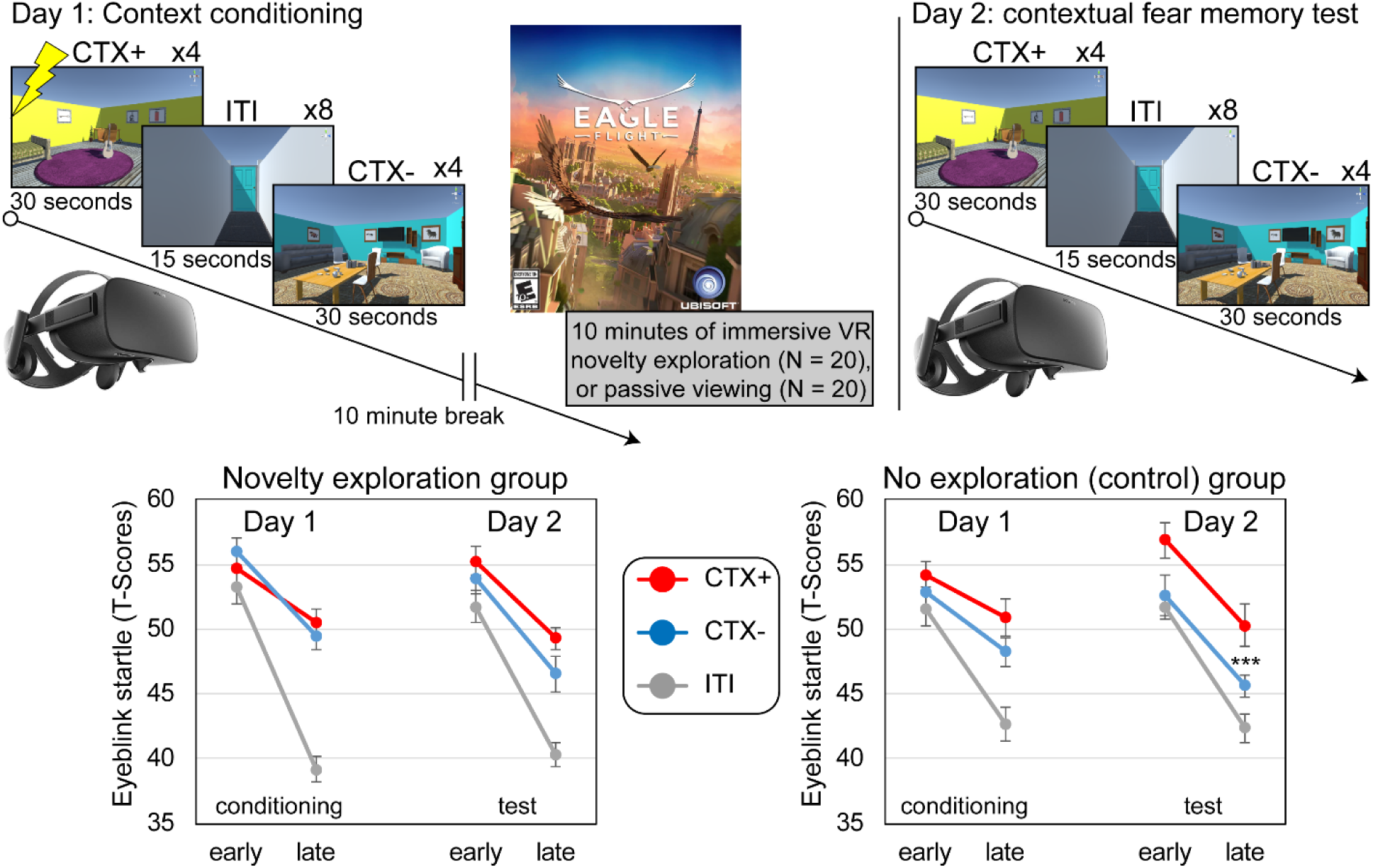
Contextual fear-conditioning paradigm and fear-potentiated startle results. a) Timeline of experimental design. After context conditioning, subjects either played a novel VR game or viewed footage of the game in 2D. Subjects returned 24 hours later for a test of contextual fear memory. b) Line plots reflecting mean startle responses in the first half (early) and second half (late) of the context-conditioning task for threat (CTX+, red), safe (CTX-, blue), and neutral contexts (ITI, grey) during Day 1 and 2 for both groups. Context conditioning resulted in stronger electromyography responses (i.e. eye-blink magnitude) to startle probes within the rooms compared to ITI during late Day 1 probes for both groups. Day 2 testing resulted in greater startle responses to late CTX+ probes compared CTX- or hallway, but with no difference between groups. Error bars = s.e.m. ***p < 0.001.

During context conditioning and test, subjects were passively guided throughout the two contexts and the hallway on a predefined path, and received an unpredictable shock in the CTX+ but not in the CTX- or the hallway. The path was slightly different each time they entered a room. Subjects passed through the CTX+ and CTX- four times each and received a total of 3 shocks in the CTX+ on Day 1. The time at which the startle probe was presented in each context, and the hallway, were pseudorandomized to prevent subjects from forming a temporal association with the shock and the rooms. The startle probe could occur randomly from 5-25 seconds after entering a context, and 5-10 seconds after entering the hallway. Subjects were told that if they paid attention they might learn an association between the rooms and the shock, but they were not instructed about the context-shock relationship and had to learn this association for themselves. Subjects were administered the Igroup presence questionnaire (IPQ) used to measure the sense of “presence” in virtual reality. There were no differences in IPQ scores for the context conditioning environments between groups (P = .95).

Following context conditioning, subjects passively watched a silent video of a train traveling through the countryside for 10 minutes, as used in some of our prior research during wait periods between experimental phases (Dunsmoor et al. 2009; Dunsmoor and LaBar 2013). Then, half the subjects played the immersive 3D video game Eagle Flight (Ubisoft) on the Oculus Rift for 10 minutes. The game involves navigating an open environment in the first person perspective of a flying eagle as you search for rewards. The other half of subjects passively watched a 10 minute video of someone else playing Eagle Flight. The IPQ scores showed much stronger sense of “presence” during Eagle Flight than during passive viewing (P < .001). The game was also rated as more “enjoyable” (P = .014), “fun” (P < .001), and “engaging” (P < .001) than the video. Two subjects did rate the VR game as making them very nauseated. Removing these subjects from the analysis did not have a meaningful impact on the results.

## Acknowledgments

The study was supported by NIH R00 MH106719 to J.E.D.

## Declaration of Interests

None.

## Contact for Resource Sharing

Further information and requests for resources should be directed to and will be fulfilled by the Lead Contact, Joseph Dunsmoor

